# Correlation between the oral microbiome and brain resting state connectivity in smokers

**DOI:** 10.1101/444612

**Authors:** Dongdong Lin, Kent E. Hutchison, Salvador Portillo, Victor Vegara, Jarrod M. Ellingson, Jingyu Liu, Kenneth S. Krauter, Amanda Carroll-Portillo, Vince D. Calhoun

## Abstract

Recent studies have shown a critical role of the gastrointestinal microbiome in brain and behavior via the complex gut–microbiome–brain axis, however, the influence of the oral microbiome in neurological processes is much less studied, especially in response to the stimuli in the oral microenvironment such as smoking. Additionally, given the complex structural and functional networks in brain system, our knowledge about the relationship between microbiome and brain function in specific brain circuits is still very limited. In this pilot work, we leveraged next generation microbial sequencing with functional neuroimaging techniques to enable the delineation of microbiome-brain network links as well as their relationship to cigarette smoking. Thirty smokers and 30 age- and sex- matched non-smokers were recruited for measuring both microbial community and brain functional networks. Statistical analyses were performed to demonstrate the influence of smoking on the abundance of the constituents within the oral microbial community and functional network connectivity among brain regions as well as the associations between microbial shifts and the brain functional network connectivity alternations. Compared to non-smokers, we found a significant decrease in beta diversity (p = 6×10^−3^) in smokers and identified several classes (Betaproteobacteria, Spirochaetia, Synergistia, and Mollicutes) as having significant alterations in microbial abundance. Taxonomic analyses demonstrate that the microbiota with altered abundance are mainly involved in pathways related to cell processes, DNA repair, immune system, and neurotransmitters signaling. One brain functional network connectivity component was identified to have a significant difference between smokers and nonsmokers (p = 0.033), mainly including connectivity between brain default network and other task-positive networks. The brain functional component was also significantly associated with some smoking related oral microbiota, suggesting a potential link between smoking-induced oral microbiome dysbiosis and brain functional connectivity, possibly through immunological and neurotransmitter signaling pathways. This work is the first attempt to link oral microbiome and brain functional networks, and provides support for future work in characterizing the role of oral microbiome in mediating smoking effects on brain activity.

## 1. Introduction

Nicotine, an addictive substance, has been reported to influence brain function and human behavior, including cognitive function and endogenous information processing networks [1]. Functional magnetic resonance imaging (fMRI) has been widely applied to delineate the interactions among diverse brain functional networks related to nicotine use or dependence, informing our understanding of the links between brain function and smoking abuse or cessation. Studies have reported negative association between the severity of nicotine dependence and dorsal anterior cingulate cortex (dACC) connectivity strength with several other regions including the ventrium and insula [2, 3]. These studies suggest the use of resting state connectivity among dACC, insula and striatum as biological measures of nicotine addiction. Consistent reduction of dACC-insula connectivity was also found in smokers who relapsed when quitting compared to those who remained abstinent [4]. Cole *et al.* [5] reported that cognitive withdrawal improvement after nicotine replacement was associated with enhanced connectivity between the executive cognitive network (ECN) and the default mode network (DMN). After acute nicotine administration, non-smokers showed reduced activity within the DMN and increased activity in extra-striate regions within the visual attention network, suggesting a shift in network activity from internal to external information processing [6]. Other evidence supports the critical role of insula, together with the ACC in influencing the dynamics between large-scale brain networks [1]. Significant lower connectivity strength between left ECN and DMN domains was found in chronic smokers compared to nonsmokers [7]. Chronic nicotine use also showed negative impact on functional network connectivity within ECN domain.

Smoking can affect oral health by altering the microbial ecosystem in the oral cavity. There are around 600 types of bacterial species inhabiting the human oral cavity, which live together in synergy [8]. Bacteria can colonize and form complex communities in the oral cavity on a range of surfaces including on the teeth, the tongue, or under the gum with each surface representing a specific microenvironment with slightly variant conditions. The oral microbiome helps to maintain oral health, but composition is sensitive to environmental disruptions including smoking cigarettes or antibiotic intake [9]. The balance of the microbial ecosystem is disturbed by these alterations, dysbiosis, resulting in diseases such as periodontitis or respiratory diseases [10–12]. Smoking can directly influence the oral microbiome and perturb oral microbial ecology through a variety of mechanisms including antibiotic effects or oxygen deprivation [13]. Evidence suggests smoking drives colonization of marginal and subgingival spaces with highly diverse biofilms resulting in a proinflammatory response from the host [14]. Investigators also found that smokers harbored more pathogenic, anaerobic microbes in the subgingival space than nonsmokers [15]. Use of 16s RNA sequencing has demonstrated a shift in the abundances of particular microbiota in smokers compared to non-smokers including an increase in pathogenic microbes associated with increased risk of oral diseases [16]. Kumar *et al.* identified an increase in periodontal pathogens belonging to the genera Fusobacterium, Cardiobacterium, Synergistes, and Selenomonas in tobacco users [14]. Wu *et al.* showed depletion in the abundance of microbes associated with oral health including from the phylum Proteobacteria and genera Capnocytophaga, Peptostreptococcus and Leptotrichia in smokers, which could potentially lead to smoking-related diseases [17].

In the past few years, many studies have shown a critical role of the gastrointestinal microbiome in brain development, function, and behavior via the complex gut microbiome–brain axis [18, 19]. It has been suggested that the communications between the microbiome and brain is bidirectional through multiple pathways including the hypothalamic-pituitary-adrenal axis (HPA), neurotransmitter pathways, immune system, and recognition of bacterial or host metabolites. Research has found that hormonal changes along the HPA axis due to neurological reactions within the brain to stress or anxiety are related to gut microbiome composition [20]. Inversely, gastrointestinal microbial perturbations have been shown to impair recognition memory and cognitive function in hippocampus [21]. Microbiota are also involved in several neurotransmitter pathways including dopaminergic, serotonergic, and glutamatergic signaling, which are well known in modulating neurogenesis and brain function [22]. Animal models have demonstrated increased levels of noradrenaline, dopamine, and serotonin in the striatum and hippocampus, and reduced expression of N-methyl-D-aspartate receptor subunits in the hippocampus, cortex, and amygdala in germ-free mice, suggesting the role of the microbiome in regulating the levels of these neurotransmitters in the brain [23]. The microbiome has also been reported to affect neurogenesis and development given its possible influence on brain-derived neurotropic factor expression in multiple brain regions. Moreover, neuroinflammation also plays a critical role in brain and behavioral abnormalities, disrupting synaptic plasticity and neurogenesis among cortical and limbic areas [24]. Certain bacteria (e.g., Bacteroidetes) are believed to stimulate neuroinflammation via increased brain-blood-barrier permeability and toll-like receptor 4 (TLR4)-mediated inflammatory pathways [25]. With such a close relationship between the gastrointestinal microbiome and brain function, researchers have identified several neurological disorders correlated to changes in gastrointestinal microbial populations including autism, major depression disorders, and neurodegenerative disorders [26–28].

While most studies focus on the influence of gut microbiome on brain signaling, the potential role of the oral microbiome in the regulation of neurological activity is much less studied. Recent work has demonstrated that oral microbial perturbations are associated with neurodegeneration (e.g., Alzheimer’s diseases, Parkinson’s disease, and glaucoma) [29–31]. Bacterial endotoxin from the oral cavity is tied to chronic, subclinical inflammation, development of neurodegeneration, and has even been localized to the brain tissue of patients suffering from Alzheimer’s disease [29]. As the second most taxonomically diverse body site, the oral microbiome consists of some bacteria that are specific to the oral cavity while also sharing microbes found within the in gastrointestinal microbiome. As such, it is not surprising that some oral and gut microbiota show concordant disease associations [32] indicating a potential connection between the two sites contributing to inflammatory diseases [33, 34]. With these linkages and close proximity to the brain, there is high potential for oral dysbiosis, similar to gastrointestinal dysbiosis, to affect brain activity. However, knowledge of the constituents and interactions of the oral microbiome is still limited, especially in relation to how the compositional changes influence neurological signaling in the context of disease.

Despite recent advances in understanding of the gut-brain axis, there is still a significant gap in knowledge as to the role of the microbiome on different regions or circuits involved in brain function. Neuroimaging is a powerful tool enabling the delineation of microbiome influences on specific brain circuitry using similar applications designed for brain functional mapping and neurological disease diagnosis [35]. Recent studies have identified the influence of changes in the gastrointestinal microbiome to activation of brain circuits related to memory and depression [36–38]. suggesting the potential of combining the microbiome and neuroimaging in studying microbiome-brain interactions. However, there have not been investigations into the link between the oral microbiome and brain function in relation to behavioral changes. In this work, we leveraged next generation 16s rRNA sequencing and resting-state fMRI techniques to explore the effects of changes in oral microbiome composition on brain function. Smoking directly influences the constitution of the oral microbiome, which allows for examination of fluctuations in brain activities that are potentially correlated with shifts of the oral microbial populations. Saliva samples and resting-state fMRI scans from 60 individuals were collected, and the associations between bacterial populations and neurological signaling (e.g., brain functional connectivity) were examined. Influence of bacterial fluctuations linked to smoking on brain functional abnormalities was demonstrated showing the potential role of oral microbiome in influencing brain functional connectivity in relation to smoking.

## 2. Materials and Method

### 2.1. Participants

Sixty subjects were used for the analyses, including 30 smokers with the level of nicotine dependence score (FTQ:Fagerstrom Tolerance Questionnaire [39]) >6 and 30 age- and sex-matched, non-smokers (FTQ score <=6). Subjects consisted of 45 males and 15 females (Fisher exact test p = 1) between the ages of 21 and 56 (37.2±10.65; p = 0.98) years. Group difference tests on age, AUDIT (the alcohol use disorders identification test [40]) score, and marijuana smoking (the number of marijuana smoking days) via two-sample t-test, and sex by Chi-square test can be seen in Table 1. Subjects with injury to the brain, brain-related medical problems, bipolar or psychotic disorders, or illicit drugs users (confirmed through urinalysis) were excluded.

**Table 1.** Demographics of subjects

|  | Smoker (n=30) | Nonsmoker (n=30) | p |
| --- | --- | --- | --- |
| Age | 37.23±9.58 | 37.17±11.78 | 0.98 |
| Sex(M/F) | 21/8 | 20/7 | 1 |
| Alcohol (AUDIT score) | 8.43±9.27 | 14.1±7.55 | 0.012 |
| Marijuana smoking | 12.27±25 | 5.57±13.11 | 0.2 |
| FTQ score | 8.87±1.57 | 0.93±1.96 | - |

### 2.2. 16S rRNA sequencing

Saliva samples were collected for 16S rRNA amplicon sequencing. Participants provided 5 ml of saliva in a sterile 50 ml conical centrifuge tube and stored in a refrigerator until the DNA was extracted. Sequencing was performed in the same laboratory using an Illumina MiSeq covering variable region V4 with primers (5’GGAGGCAGCAGTRRGGAAT-3’ and 5’-CTACCRGGGTATTAAT −3’). Raw sequence data were demultiplexed, followed by quality control by applying the pipeline in DADA2 [41]. generating a number of unique sequences, similar to operational taxonomic unit (OTU) derived by sequences clustering with 100% identity accuracy in the previous pipeline [42]. Each sequence was trimmed to have a length of 150 base pairs and was aligned by the mafft tool to build a phylogenetic tree [43]. A classifier for taxonomy assignment was trained based on sequences and taxonomic results from Greengenes database (http://greengenes.lbl.gov) with a 99% similarity. The classifier was then applied to the identified sequences for taxonomic assignment. Assigned sequences were further agglomerated at the genus, family, and class levels for taxonomic analyses. All processing scripts were implemented on a QIIME2 (https://qiime2.org/) platform.

### 2.3. Resting state fMRI imaging

Fifty-six participants also had resting state functional MRI (rsfMRI) collected on a 3 T Siemens TIM Trio (Erlangen, Germany) scanner. Images were acquired with an echo-planar imaging (EPI) sequence (TR=2000 ms, TE=29 ms, flip angle=75°) with a 12-channel head coil. Each volume consisted of 33 axial slices (64×64 matrix, 3.75×3.75 mm^2^, 3.5 mm thickness, 1 mm gap). Image preprocessing was performed as previously described [44]. Briefly, this included slice-timing correction, realignment, co-registration and spatial normalization. By transforming the images to the Montreal Neurological Institute (MNI) standard space, we kept those with the root mean square of head movement not exceeding 3 standard deviations, despiked time courses[45]. and smoothed images using a FWHM Gaussian kernel of size 6 mm. The data were then analyzed by group independent component analysis (GICA) [46] with 120 and 100 components for the first and second decomposition levels respectively [46, 47]. Thirty-nine out of the 100 components were selected with low noise and free of major artifacts. The spatial map of each selected component was z-transformed, and voxelwise one-sample t-test statistics were thresholded to identify the main brain areas. The time course corresponding to each component was filtered using a band-pass filter 0.01-0.15 Hz. Finally, resting state functional network connectivity (FNC) matrices were calculated for each subject based on the correlation coefficients between the time courses of all possible pairs formed with the 39 chosen components.

### 2.4. Analysis of oral microbiome

We tested the overall microbiota composition difference between smoking and non-smoking groups by comparing cross-sample distance. Raw read counts were first rarefied at 2020 sequences/sample. Weighted and unweighted UniFrac distances and Bray-Curtis distance were assessed by the R package ‘vegan’ [48] and tested for group difference by applying permutational MANCOVA (‘Adnois’ function in vegan package) controlling for age, sex, alcohol AUDIT score, and marijuana smoking score. Principal coordinate analysis (PCoA) plots were generated based on the first two principle coordinates from each type of distance matrix to compare the dissimilarity among samples in the subspace spanned by the first two dimensions.

The OTU table of raw read counts was normalized to the table of relative abundances at different taxa levels. The taxa present in less than 20% of subjects were filtered out, resulting in 163 OTUs, 73 genera, and 20 classes. Each taxon was tested for the relative abundance difference between smoking and non-smoking groups by Wilcox ranked sum test. Those taxa with significant group difference, were further tested controlling for age, sex, alcohol and marijuana smoking by *‘Zig’* function in the *MetagenomeSeq* package [49]. The *‘Zig’* method has demonstrated advantage in microbiome data analysis by modeling raw counts using multivariate Gaussian distribution and taking into account zero abundance in a large proportion of subjects for each taxon.

### 2.5. Analysis of resting state fMRI imaging

Each element in the FNC matrix indicates the correlation between any two functional networks within the brain. We transformed the FNC matrix of each subject to the vector and concatenated all FNC matrices across all subjects to form the matrix ***C*** with a dimension of *n* (subjects) by *m* (FNC links). To reduce the dimension of large number of FNC links, we applied the ICA algorithm to decompose matrix ***C*** into the multiplication of two full rank matrices by ***C*** = ***AS***, where matrix ***S*** contains independent FNC components with similar cross-individual pattern and matrix ***A*** is the loading matrix which shows the presentation of each component on the subjects. The ICASSO algorithm [50] followed by best run selection was applied to obtain four reliable FNC components and each corresponding loading was tested for group difference by two-sample t-test. For the component with significant group difference, multiple regression model was further applied by controlling for covariates.

### 2.6. Linking microbiota with FNC

After obtaining significant FNC component and taxa with the above analyses, we further tested the association between FNC component and taxa using the *‘Zig’* function with the following model design:

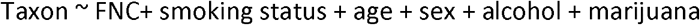

We tested taxa from OTU, genus, and class levels, and reported the significant associations with nominal p<0.05. Multiple comparisons in all tests were corrected by false discovery rate (FDR) method.

### 2.7. Functional analysis of predicted metagenomes

Metagenome content in the samples was inferred from 16S rRNA microbial data, normalized by copy number count to account for the differences of the number of 16S rRNA copies between taxa, and then functional metabolic pathways were predicted based on the Kyoto Encyclopedia of Gene and Genomes (KEGG) catalog [51]. using Phylogenetic Investigation of Communities by Reconstruction of Unobserved States (PICRUSt) [52]. Analyses revealed 328 metabolism pathways at level 3 were predicted. Of these, 66 pathways were removed due to presence in less than 10% of samples. Group difference of each metabolism pathway between smokers and non-smokers was tested using Welch’s t-test using the Statistical Analysis of Metagenomic Profiles (STAMP) software [53]. Multiple comparisons were corrected by FDR with cut-off set as 0.15 for significance. For the KEGG pathways and OTUs or genera significantly associated with smoking status, we further used Spearman’s rank correlation to examine their relationship.

## 3. Results

### 3.1. Overall microbiome composition between smoking groups

To determine whether overall microbiome composition differed between smokers and nonsmokers, we performed principal coordinate analysis on unweighted UniFrac, weighted UniFrac and Bray-Curtis phylogenetic distances. Without controlling for covariates, we found significant group difference in unweighted UniFrac distance (p= 6×10^−3^) and Bray-Curtis distance (p= 0.025) under 9999 times permutation. The differences were still significant (p= 4×10^−3^ and 0.027, respectively) after controlling covariates (age, sex, alcohol score and marijuana smoking). No significant differences were found in weighted UniFrac (p= 0.2).

**Figure 1.**
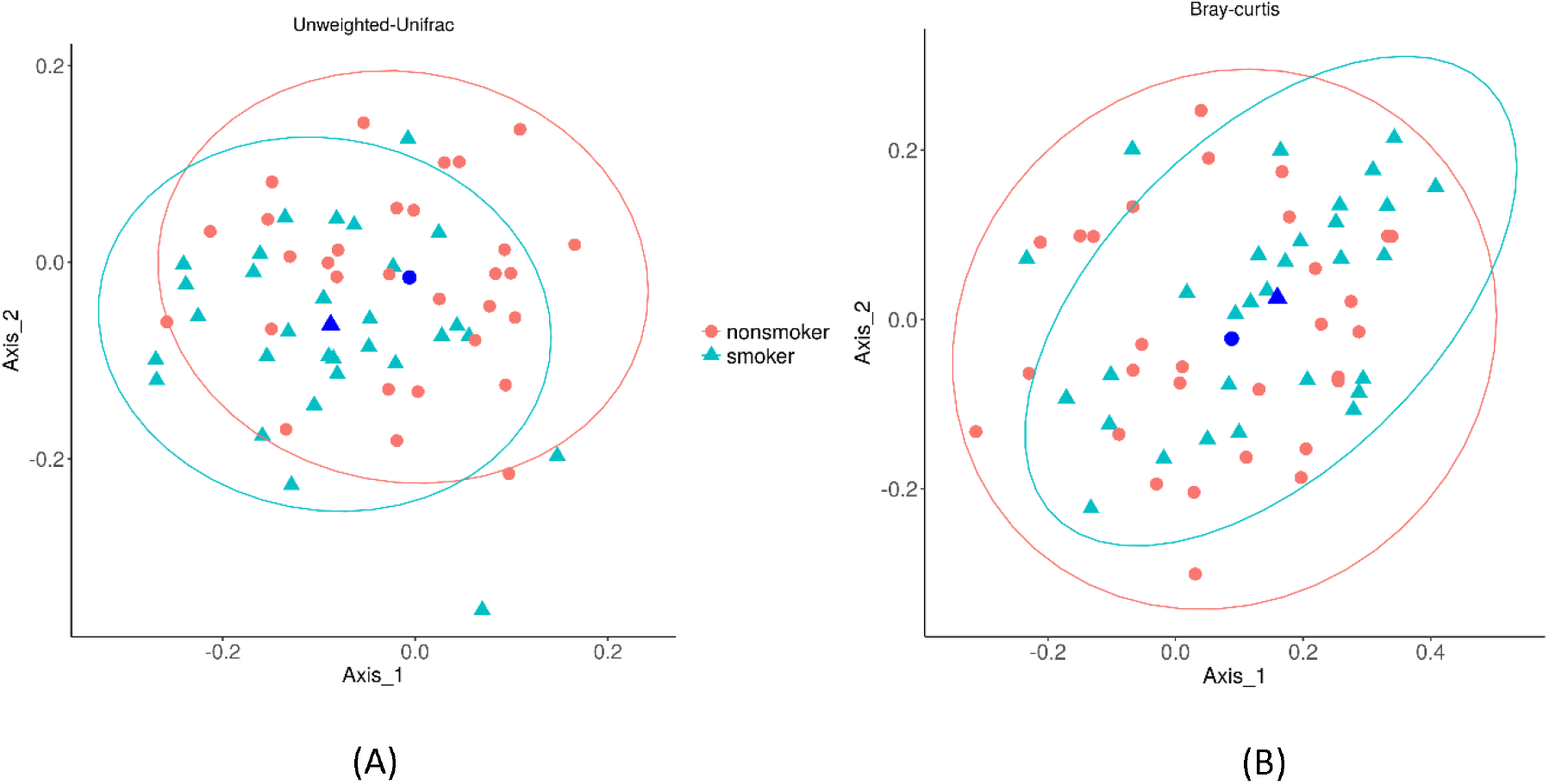
PCoA analysis of microbial composition between smokers and non-smokers. The microbial composition was evaluated based on (A) Unweighted UniFrac distance and (B) Bray-Curtis distance, respectively. Dark blue point indicates the center of eclipse.

### 3.2. Taxonomic analysis between smokers and nonsmokers

To determine the compositional differences in the salivary microbiota of smokers and nonsmokers, we examined the relative abundance of taxa at OTU, genus, and class levels. For each taxon, we tested their group difference using non-parametric rank sum test. For taxa with significant group difference (FDR≤ 0.05), a multivariate test was applied controlling for covariates (age, sex, alcohol score and marijuana smoking). Fig.S1 and Table 2 show the relative abundance and log fold change (logFC=log2(smokers/nonsmokers)) of significant taxa by multivariate test at the OTU, genus, and class levels. We found that class *Betaproteobacteria* significantly differed in smokers with a clear depletion as compared to non-smokers (logFC= -0.35, p= 3.2×10^−2^). Within this class, it was genera *Lautropia* (logFC=-1.99, p= 7.4×10^−3^) and *Neisseria* (logFC= -1.16, p= 8.5×10^−3^) that were specifically reduced in smokers. Other genera displaying significant differences between smokers and non-smokers included *Treponema* (class Spirochaetes), *TG5* (class Synergistia), and *Mycoplasma* (class Mollicutes), which all had significant enrichment in smokers. Additionally, within the smoking population, there was a significant increase in relative abundance of both genus *Bacteroides* and the family *Mogibacteriaceae* (logFC= 2.24, p= 7.9×10^−4^; logFC= 2.92, p= 3.1×10^−4^, respectively) but not seen at the class level.

**Table 2:**
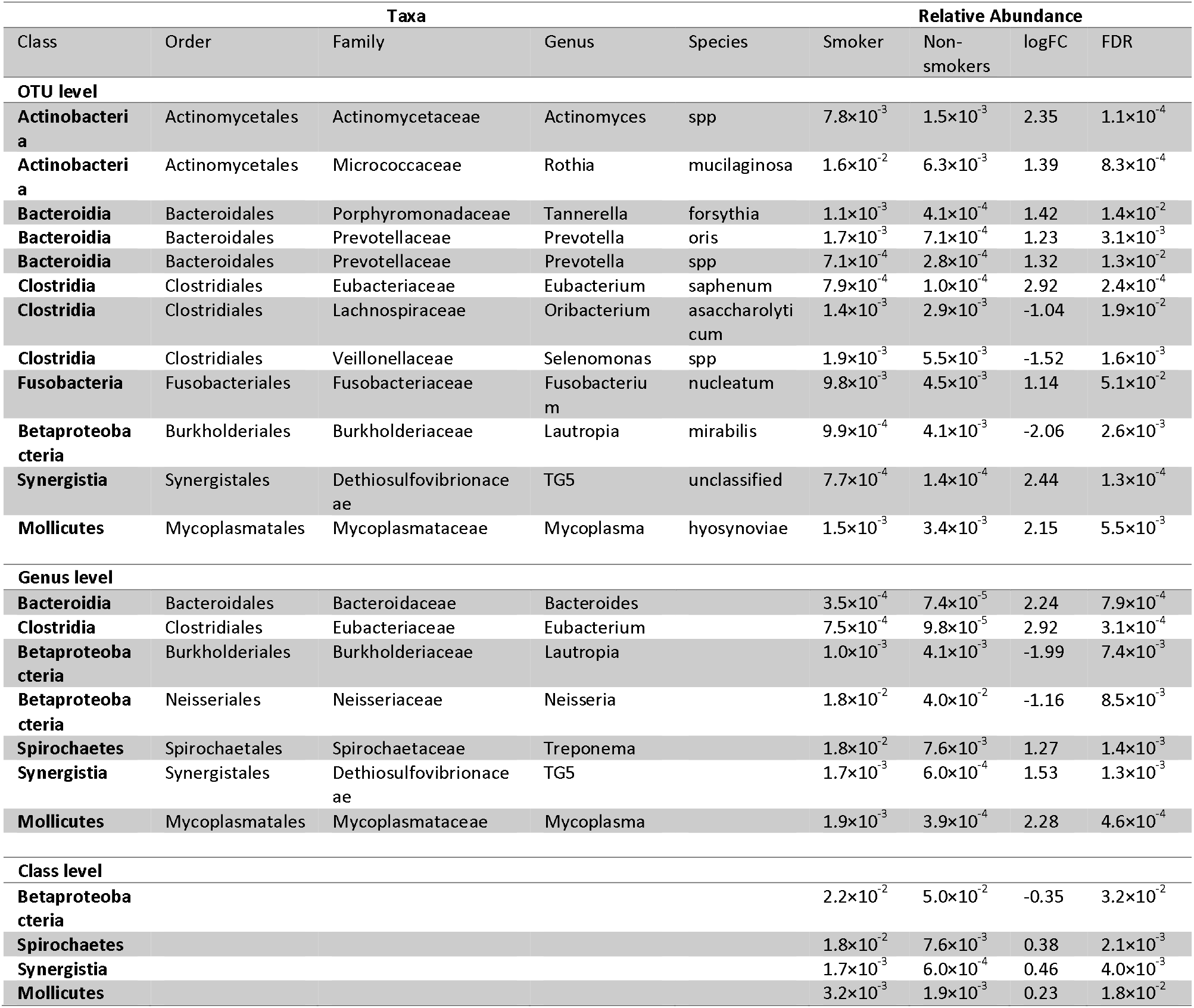
List of taxa with significant difference between smoking and non-smoking groups at species, genus and class levels.

Lower-level analyses on OTU identified 12 OTUs from 7 classes showing significant difference between smoking and non-smoking groups. Besides the classes identified at genus level, we additionally found 2 OTUs from genus *Actinomyces* and *Rothia* in class Actinobacteria with higher abundance in smokers compared to non-smokers. The abundance of three OTUs from genera *Tannerella* and *Prevotella* (in class Bacteroidia) and one from genus *Fusobacterium* were also significantly increased in smokers. On the contrary, 2 OTUs from genera *Oribacterium* and *Selenomonas* were depleted in smokers.

### 3.3. Differential FNC component between groups

Among four rsfMRI components derived by applying ICA on the FNC matrix, there was one component with the corresponding loading vector showing significant difference between smoking and non-smoking groups (nominal p= 0.033), as shown in Fig. 2A. The group difference remained significant after controlling for covariates (p= 0.04) suggesting higher loading in smokers compared to nonsmokers for the FNC component. All of the top 13 FNCs with absolute (z-scored weights)> 2.5 in the component had negative weights (Fig. 2B), which indicates that those FNCs contributed in an opposite way to the component that they demonstrated significant reduction in smoking group. Fig. 2C shows the connectivity among functional networks from the top contributing FNCs in the component. The brain regions of those connected functional networks were plotted in Fig. 2D. Altered connectivity was mainly between DMN and visual network (VIS), salience network (SAL), and cognitive controls network (ECN), as well as between precunes (PRE) network and VIS. As listed in Table S1, the DMN network included several functional regions including anterior cingulate cortex (ACC), left angular gyrus and posterior cingulate cortex (PCC) with some precuneus overlap. The VIS group was composed of right fusiform/lingual gyrus, left middle occipital gyrus and right inferior occipital gyrus. The FNCs between DMN and other task-positive networks (VIS domain, inferior frontal gyrus (IFG) within the ECN domain, right supramarginal gyrus within the SAL domain, precuneus from PRE domain and supplementary motor area from SEN domain) were all negative, indicating stronger anti-correlation between DMN and those task-positive networks in smokers.

**Figure 2.**
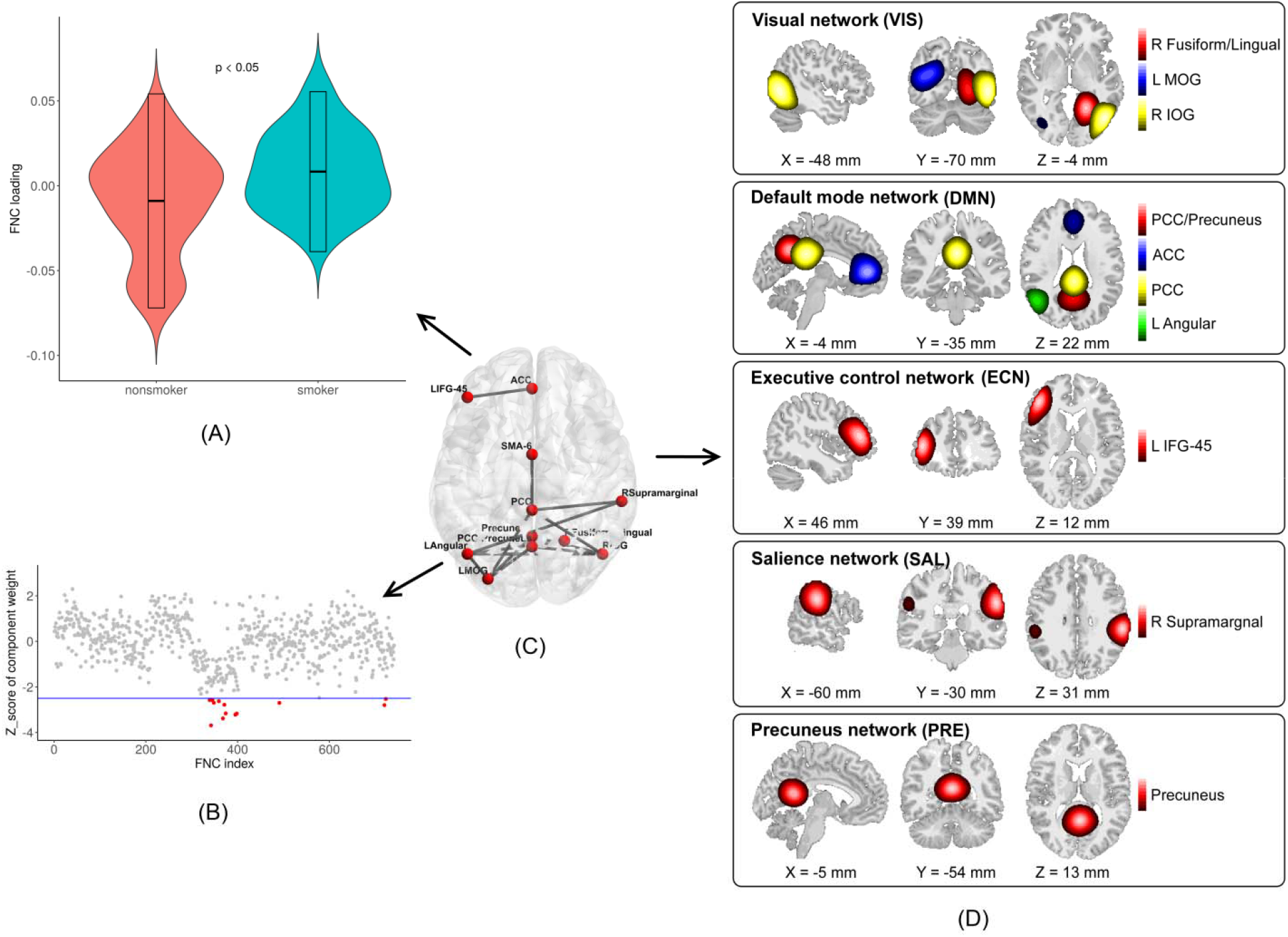
The loading and spatial mapping of identified FNC component. (A) The loading of identified FNC component in smokers compared to nonsmokers; (B) Top FNCs with z-scored weights absolute (z-score)> 2.5 in the component; (C) The top contributing connectivity among the functional networks from the selected component; (D) Brain regions of those functional networks involved the top FNCs.

### 3.4. Association test between FNC component and microbiome

After identifying both taxa and FNC component showing significant group difference, we further tested for associations between relative abundance of each taxon and the loading of FNC component. Table 3 lists significant associations between taxa and FNC component. The abundance of *Treponema* (class *Spirochaetes*),*TG5* (class *Synergistia*) and *Eubacterium* (class *Clostridia*)were positively associated with the FNC loading of the component while *Neisseria* (class *Betaproteobacteria*) demonstrated an opposite relationship. Genus *Bacteroides* (class *Bacteroidia*) also had a marginal association, suggesting higher abundance of *Bacteroides* related to higher FNC loading and thereby lower connectivity strength of top FNCs in this component. Lower-level analysis further identified that the abundance of species from genus *Actinomyces* (class *Actinobacteria*) showed significant positive relationship with FNC loading only in smokers (logFC= 2.36, p= 4.9×10^−7^), while genera *Prevotella* (class *Bacteroidia*) and *Rothia* (class *Actinobacteria*) were significantly increased along with lower top FNCs strength in smokers.

**Table 3.** The associations between smoking-related taxa and FNC component

| Taxa |  |  |  |  | Association with FNC component |  |  |
| --- | --- | --- | --- | --- | --- | --- | --- |
| Class | Order | Family | Genus | Species | log FC | Statistic | FDR |
| OTU level |  |  |  |  |  |  |  |
| Actinobacteria | Actinomycetales | Actinomycetaceae | Actinomyces | mucilaginosa | 2.35 | 120.9 | 4.9×10<sup>-7+</sup> |
| Actinobacteria | Actinomycetales | Micrococcaceae | Rothia | forsythia | 1.39 | 19.8 | 6.2×10<sup>-2</sup> |
| Bacteroidia | Bacteroidales | Prevotellaceae | Prevotella | oris | 1.23 | 43.2 | 5.8×10<sup>-6</sup> |
| Clostridia | Clostridiales | Eubacteriaceae | Eubacterium | saphenum | 2.92 | 34.0 | 1.1×10<sup>-4</sup> |
| Genus level |  |  |  |  |  |  |  |
| Bacteroidia | Bacteroidales | Bacteroidaceae | Bacteroides |  | 2.24 | 15.12 | 6.2×10<sup>-2</sup> |
| Clostridia | Clostridiales | Eubacteriaceae | Eubacterium |  | 2.92 | 35.09 | 7.1×10<sup>-5</sup> |
| Betaproteobacteria | Neisseriales | Neisseriaceae | Neisseria |  | -1.16 | -20.18 | 3.7×10<sup>-2</sup> |
| Spirochaetes | Spirochaetales | Spirochaetaceae | Treponema |  | 1.27 | 18.78 | 5.0×10<sup>-2</sup> |
| Synergistia | Synergistales | Dethiosulfovibrionaceae | TG5 |  | 1.53 | 13.04 | 3.7×10<sup>-2</sup> |
| Class level |  |  |  |  |  |  |  |
| Betaproteobacteria |  |  |  |  | -0.35 | -20.15 | 3.7×10<sup>-2</sup> |
| Synergistia |  |  |  |  | 0.46 | 14.80 | 2.9×10<sup>-2</sup> |
| Mollicutes |  |  |  |  | 0.23 | 18.76 | 1.9×10<sup>-2</sup> |

+: the significance was shown in smoking group only.

### 3.5. Functional metabolism pathway prediction

Among the 262 KEGG pathways predicted for microbial function, we identified 23 pathways showing significant difference in abundance between smokers and non-smokers after correcting for multiple tests by FDR with a threshold 0.15, as shown in Fig.3. These pathways with abundance significantly altered in smokers mainly involved metabolism and genetic information processing. Enriched metabolic pathways included those involved with metabolism (cofactors and vitamins, terpenoids and polyketides, amino acids, nucleotides, and glycans). Depleted pathways were involved with energy and lipid metabolism, membrane transport, and xenobiotics biodegradation (i.e., drug metabolism-cytochrome P450). In addition, genetic information processing pathways (i.e., proteasome, protein export, nucleotide excision repair, DNA repair and recombination proteins, and ubiquitin system) were also significantly enriched in smokers whereas other pathways related to diseases (i.e., immune disease, neurodegenerative disease), nervous system, and circulatory system were also associated. Further examination of the correlations between microbiota and the functional pathways demonstrated that alterations in the abundance of microbiota were highly correlated to these pathways (mean absolute value of correlation 0.19~0.29 in different taxonomic levels), especially for the genera *Lautropia* and *Neisseria* from class Betaproteobacteria as shown in Fig. S2.

**Figure 3.**
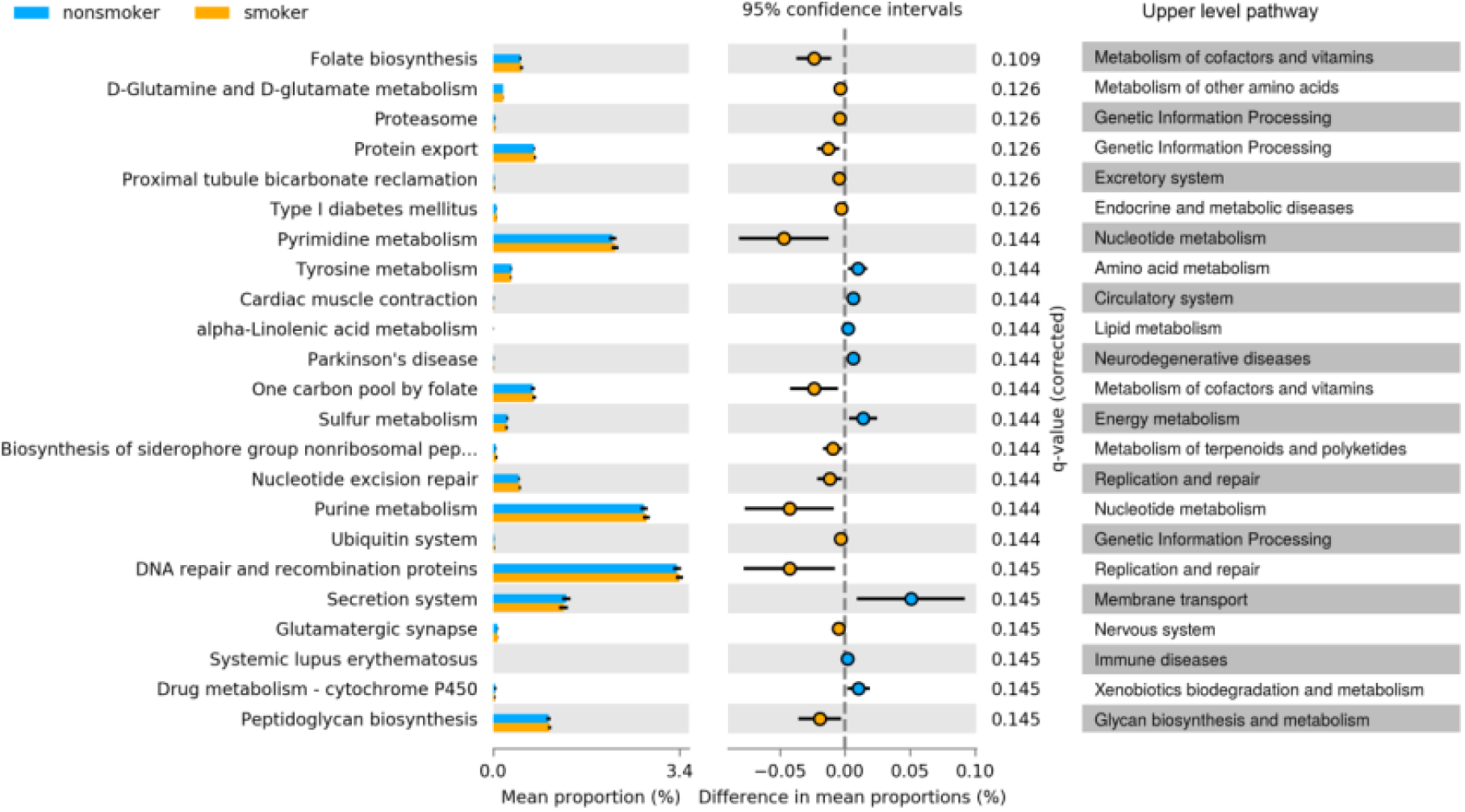
Pathway enrichment analysis based on the predicted metagenomics. The 23 out of 262 KEGG functional pathways show significant changes in abundance between smokers and non-smokers (FDR < 0. 15). Those pathways were predicted from 16S rRNA microbiome sequencing using the PICRUSt algorithm. Mean proportion (colored bar) indicates the relative abundance of the pathway in each group. The difference of mean proportions between groups as well as the 95% confidence interval indicates the effect size of relative abundance change for each pathway.

## 4. Discussion and Conclusions

In this work, we specifically set out to determine if correlations existed between shifts in the oral microbial population and changes in brain signaling networks due to smoking. To achieve this, we used 16s rRNA sequencing to characterize the microbial composition in the saliva of participants (smokers versus nonsmokers), and rsfMRI to measure brain functional activity in these same participants. Data delineating microbial shifts was consistent with previously reported findings for changes in the oral cavity due to smoking. Likewise, changes in brain functional activity also matched with previous results found due to smoking. When correlative analyses were performed on with these data sets, some oral microbial populations were found to have significant correlation with particular neurological signaling networks. While the influence of the gastrointestinal microbiome on neurological activity has been an area of intense study, this study suggests that the oral microbiome also influences neurological signaling and may provide new therapeutic opportunities for treatment of neurological disorders.

As stated, results from 16s rRNA sequencing were consistent with previously reported results for changes in the oral microbiome due to smoking. We found significant changes in microbial composition in both unweighted UniFrac and Bray-Curtis distances with smaller variation within smokers than nonsmokers, showing less microbial diversity in the salivary microbiome of smokers. Taxonomic analyses identified drastic abundance changes on multiple taxa at different levels. Gram-negative bacteria from the genera *Loutropio* and *Neisseria* from the class *Betaproteobacteria* showed depletion in the smokers, in line with previous [14, 16]. Several in vitro studies have also demonstrated the strong inhibitory effect of smoking in *Neisseria* growth [54]. Other genera were enriched in smokers including *Bacteroides* [55]. *Treponema* [14]. *TG5* and *Mycoplasma* [55]. especially *Mycoplasma* may be synergize with smoking to produce the pro-inflammatory effects [56]. In addition, low-level analysis identified some species significantly enriched in smokers. *Actinomyces spp*. and *Rothia mucilaginosa* from the gram-negative class *Actinobacteria* were increased in abundance in smokers, in line with previous large scale oral microbiome study [17]. *Tannerella forsythia* has been reported to enrich in subgingival plaque of current smokers and demonstrated potential risk as pathogen to induce periodontal disease [57]. *Prevotella oris* and *Prevotella spp*. from *Bacteroidales* are coaggregated with Porphyromonas gingivalis [58] which is also a critical periodontitis pathogen and is highly promoted during the infection by smoking [59]. Most of these enriched microbiota are anaerobes compared to aerobic *Neisseria*, consistent with the finding of higher abundance of anaerobes in subgingival plaque samples of smokers, suggesting the depleting of oxygen in oral cavity induced by smoking [15].

KEGG pathway analysis identified several metabolic pathways involved in functional changes during smoking. This is perhaps not surprising as cigarette smoke has been reported to be highly associated with DNA damage, lipid peroxidation and antioxidant impairment, and protein modification and misfolding, thereby inducing severe cellular damage [60, 61]. These influences may affect the oral microbiome community with its direct proximity to toxins from cigarette smoking. We found significant enrichment of metabolic pathways involving the proteasome, protein export and the ubiquitin system. All of them are related to protein degradation and recycling in the cell, which is essential for cellular processes such as proliferation, signaling, and immune responses [62]. The up-regulation of these pathways indicates the role of smoking in disrupting protein modification and cellular processes of the microbial constituents within the oral cavity. Other pathways related to DNA repair and replication-including folate biosynthesis- were also significantly activated in the oral microbiome of smokers. The involvement of proteasome function and DNA repair pathways enhanced in the smoking population may be due to increased cellular dysfunction and DNA damage induced by smoking. Additionally, enrichment was also found in pathways related to small amino acid production such as glutamate and glutamine, glutamatergic synapse and tyrosine metabolism, which are related to neurotransmitter release and potentially interact with nervous system in changing neuronal activity of smokers such as addiction and craving [63, 64]. On the contrary, some metabolisms are significantly depleted in smokers such as metabolisms of lipid energy (i.e., alpha-linolenic and sulfur), and xenobiotics biodegradation (i.e., drug metabolism-cytochrome P450), which is in line with previous studies [17, 65].

Neuroimaging analysis identified one smoking-related FNC component involved in the connectivity between DMN and other task-positive networks from VIS, SAL, ECN and PRE domains, which is consistent with our previous study [44]. DMN is mostly related to self-referential and episodic memory processing, which is down-regulated in task performance [66]. The other networks are activated corresponding to different tasks such as visual, cognitive control, attention and moment-to-moment information processing, namely ‘task-positive’ regions [67]. Functional MRI studies in both task performance and resting state have reported tight coupling between DMN and other task positive networks with negative correlations (anti-correlation) [67–69]. In our work, we found similar anticorrelations between DMN and the other domains (e.g., VIS, SAL and ECN) with reduced connectivity (i.e., increased negative coupling) in smokers when compared to nonsmokers. Reduced connectivity within and between DMN and ECN networks were also reported in chronic smokers compared to nonsmokers, showing larger decreases of connectivity with heavier nicotine use [7]. Additional studies found increased coupling among medial orbital prefrontal cortex, the dorsal medial PFC, striatum, and visual cortex over the course of 1 h acute abstinence, which is consistent with our findings [70]. The activation of these regions, which relate to reward system and also fall in the DMN domain, indicates the relation of DMN with cigarette craving. Our results combined with the above findings suggest dynamic modulation in functional coupling between DMN and task-positive networks when subjects smoke or go through withdrawal as compared to nonsmokers.

By correlating changes in oral microbial abundance with the smoking-related brain FNC component, we identified several microbiota related to brain function. *Prevotella oris* (class *Bacteroidia*), a common gram-negative, anaerobic bacterium of the normal oral flora has been associated with the development of brain absences and other neurological syndromes (i.e., Lemierre’s syndrome) through production of IgA proteases to promote virulence and initiate an immune response [71, 72]. Another member of the class *Bacteroidia, genus Bacteroides*, has the ability to produce complex, pro-inflammatory neurotoxins that may induce inflammation in oral cavity and further contribute to development of inflammation in the brain, increasing brain-blood-barrier permeability through the circulatory system [73]. *Genus Neisseria*, including *species Neisseria meningitidis*, stimulates the immune system through a variety of mechanisms (e.g., the production of lipopolysaccharide endotoxin) and invades the neurological nervous system during infection [74]. Similarly, *Treponema* infects the brain via branches of the trigeminal nerve [75]. All of these bacteria, whose populations are influenced by smoking, affect the immune system and are capable of influencing neurological processes through either direct or indirect means. In this study, we demonstrate that these bacteria have significant associations with brain function alternated by smoking, suggesting potential pathways (i.e., inflammatory pathways) exist for members of the oral microbiome to influence neurological signaling in the brain, similar to how gut microbiome-brain interactions occur [76]. Along the gut-brain axis, neurotransmitter signaling pathways play important roles in bidirectional modulation. Our functional pathway prediction analysis identified enrichment in several neurotransmitter-related pathways among oral microbiota such as glutamate-glutamine and glutamatergic synapse. Production of neurotransmitters from these pathways (i.e., glutamate and glutamine) is stimulated by smoking and they are highly involved in reward circuit neural functions for smoking dependence, or craving after smoking withdraw [77, 78]. The high correlations between some FNC-related microbiota and these neurotransmitters signaling pathways demonstrates the potential of these specific microbiota together with other oral microbiota to influence brain function through neurotransmitter signaling pathways.

This study explores the association between fluctuations within the oral microbiome and the brain functional network in smokers. While some associations were identified here, sampling of a larger population would strengthen these findings. Additionally, although we tried to control for alcohol consumption and marijuana smoking score, their complex interactions with cigarette smoking with respect to the oral microbiome community and brain function may still confound our results. As such, follow on studies should employ stricter criteria for selection of the smoking and non-smoking control group are suggested. Despite these limitations, this study represents the first evidence of correlation between population shifts within the oral microbiome and changes in neurological signaling. As the oral cavity is an easily accessible environment, as compared to the gastrointestinal tract, further study offers opportunity for development of novel therapeutics for neurological syndromes.

## Acknowledgements

This work was supported by the National Institutes of Health [grant numbers P20GM103472 and R01EB005846], and National Science Foundation, grant number: 1539067.

The authors declare that they have no competing interests.

